# *C. elegans* LIN-66 mediates EIF-3.G-dependent protein translation via a cold-shock domain

**DOI:** 10.1101/2024.02.20.581241

**Authors:** Stephen M. Blazie, Daniel Fortunati, Yan Zhao, Yishi Jin

## Abstract

Protein translation initiation is a complex and conserved process involving many proteins acting in concert. The eukaryotic initiation factor 3 (eIF3) complex is essential for the assembly of the pre-initiation complex that scans and positions mRNA at the initiation codon. eIF3 complex consists of 13 subunits. In addition to their essential roles in general translation initiation, emerging studies suggest that individual subunits of eIF3 can provide specialized functions in response to specific stimuli. We have previously reported that a gain-of-function (gf) mutation in the G subunit of *C. elegans* eIF3 complex, *eif-3.g(gf),* selectively modulates protein translation in the ventral cord cholinergic motor neurons. Here, through unbiased genetic suppressor screening, we have identified the *lin-66* gene that mediates the *eif-3.g*(*gf*)-dependent protein translation in the motor neurons. LIN-66 is previously reported to be a nematode-specific protein composed of largely low complexity amino acid sequences with unknown functional domains. We combined bioinformatic analysis with *in vivo* functional dissection and identified a cold-shock domain in LIN-66 to be critical for its function. In the cholinergic motor neurons, LIN-66 shows somatic cytoplasmic localization and close association with EIF-3.G. The low complexity amino acid sequences of LIN-66 modulate its subcellular pattern. Cold-shock-domains are known to interact with RNA and have broad functions in RNA metabolism and protein translation. We propose that LIN-66 mediates stimuli-dependent protein translation by facilitating the interaction of mRNAs with EIF-3.G.

## Introduction

Protein translation initiation in eukaryotic cells is highly regulated and involves coordinated actions of multiple protein complexes, known as eIF1-6 (Hinnebusch and Lorsch, 2012). The eIF complexes work in concert to orchestrate orderly assembly of ribosome subunits with mRNAs and initiator tRNA. eIF3 is the largest translation initiation protein complex, consisting of 13 subunits, and recruits the small ribosomal subunit and mRNAs to form the 43S pre-initiation complex, which scans the 5’ untranslated region of mRNAs to locate the initiation codon (Cate, 2017). Consistent with their critical roles in protein translation, knockout or knockdown of many subunits of eIF3 generally results in inviable cells or organisms (Hanachi et al., 1999; Kamath et al., 2003; Zhuo et al., 2023). However, recent studies have suggested that specific subunits of eIF3 can provide precise temporal and spatial regulation of protein translation in cell proliferation, development, and stress response (Lee et al., 2015; Lee et al., 2016). Emerging mechanistic studies suggest that such eIF3 subunit-specific functions can be achieved through stimuli-induced protein phosphorylation and/or specific protein interaction partners (Lamper et al., 2020).

High resolution cryo-electron microscopy and X-ray structures of the mammalian and yeast 43S pre-initiation complex have been reported (des Georges et al., 2015; Erzberger et al., 2014; Hashem et al., 2013; Querol-Audi et al., 2013). Eight subunits of eIF3 form a core module and 5 subunits are located at the periphery of the 43S complex. The g subunit of eIF3 resides in the periphery of the complex, binds the core subunit eIF3.I through its N-terminal region and interacts with RNA via an RRM at the C-terminus (Hanachi et al., 1999). The middle region of eIF3g has a Zinc-Finger (ZF) that is not yet visible in any structures. Dysregulation of eIF3g has been implicated in human diseases. For example, altered expression of eIF3g is associated with narcolepsy (Holm et al., 2015) and also observed in an animal model for autism (Hornberg et al., 2020). In Drosophila sensory neurons knockdown of eIF3g impairs dendrite pruning (Rode et al., 2018).

We have previously reported a role of the *C. elegans* g subunit of eIF3, eif-3.G, in shaping the neuronal protein landscape (Blazie et al., 2021). A gain-of-function mutation in a neuronal acetylcholine receptor, *acr-2(gf),* causes hyperexcitation of the ventral cord cholinergic motor neurons, and the *acr-2(gf)* animals exhibit spontaneous whole-body convulsions (Jospin et al., 2009). We identified a missense mutation in the ZF of EIF-3.G in a genetic screen for behavioral suppression of *acr-2(gf)* mutants. Null mutants of *eif-3.G* are arrested in larval development. However, *eif-3.G* mutants that carry either the missense mutation or a small deletion of the ZF, designated *eif-3.G(gf),* behave indistinguishably from wild type in movement and health, but act cell-autonomously in the cholinergic motor neurons to ameliorate convulsion of *acr-2(gf)*. Using neuron-specific mRNA cross-linking immunoprecipitation and deep-sequencing (CLIP-seq), we have defined a mechanism by which EIF-3.G(GF) selectively modulates translation of a set of mRNAs with GC-rich 5′ UTRs (untranslated regions) in the hyperexcited cholinergic motor neurons (Blazie et al., 2021).

Here, we report that the function of EIF-3.G(gf) in tuning neuronal protein translation involves the LIN-66 protein. We performed a genetic screen to search for factors that reverted the behavioral suppression of *eif-3.G(gf)* on the convulsion of *acr-2(gf)* animals. We identified a loss of function mutation in *lin-66*, which was previously reported to be conserved in nematode species, with unknown functional domains (Morita and Han, 2006). Through informatic analysis of homologous protein structures, we identified that LIN-66 has a cold-shock domain embedded within low-complexity protein sequences. LIN-66 localizes to somatic cytoplasm, closely associated with EIF-3.G. The localization pattern of LIN-66 depends on the low-complexity sequences. By functional dissection, we show that the cold-shock domain is critical for LIN-66 function. Our data uncovers a previous unknown function of LIN-66 and provides *in vivo* insight to how EIF-3.G modulates neuron-type specific protein translation.

## Results

### Loss of function in *lin-66* suppresses the gain-of-function effect of *eif-3.G* on convulsion of *acr-2(gf)*

We aimed to identify genes that function in *eif-3.G(gf)-*dependent protein translation in hyperactivated cholinergic motor neurons of *acr-2(gf)* animals. We reasoned that genetic mutations reversing the suppression of *acr-2(gf)* convulsion by *eif-3.G(gf)* would likely reveal clues to such genes. We mutagenized *eif-3.G(gf); acr-2(gf)* double mutants, which exhibited wild-type like locomotion. We screened F2 progeny for animals that displayed convulsions resembling *acr-2(gf)* single mutants (see Methods). Following outcrossing analysis, we were able to re-isolate one mutation, *ju1661,* that exhibited specific suppression on *eif-3.G(gf)* in the context of *acr-2(gf)*. We then combined whole-genome sequencing analysis with single-nucleotide-polymorphism (SNP) mapping of recombinants to localize *ju1661* to the right arm of chromosome IV. Examination of candidate SNPs that likely disrupt a gene’s function revealed a point mutation changing the splice-acceptor site of the third exon of the *lin-66* gene (Figure 1A).

**Figure 1.**
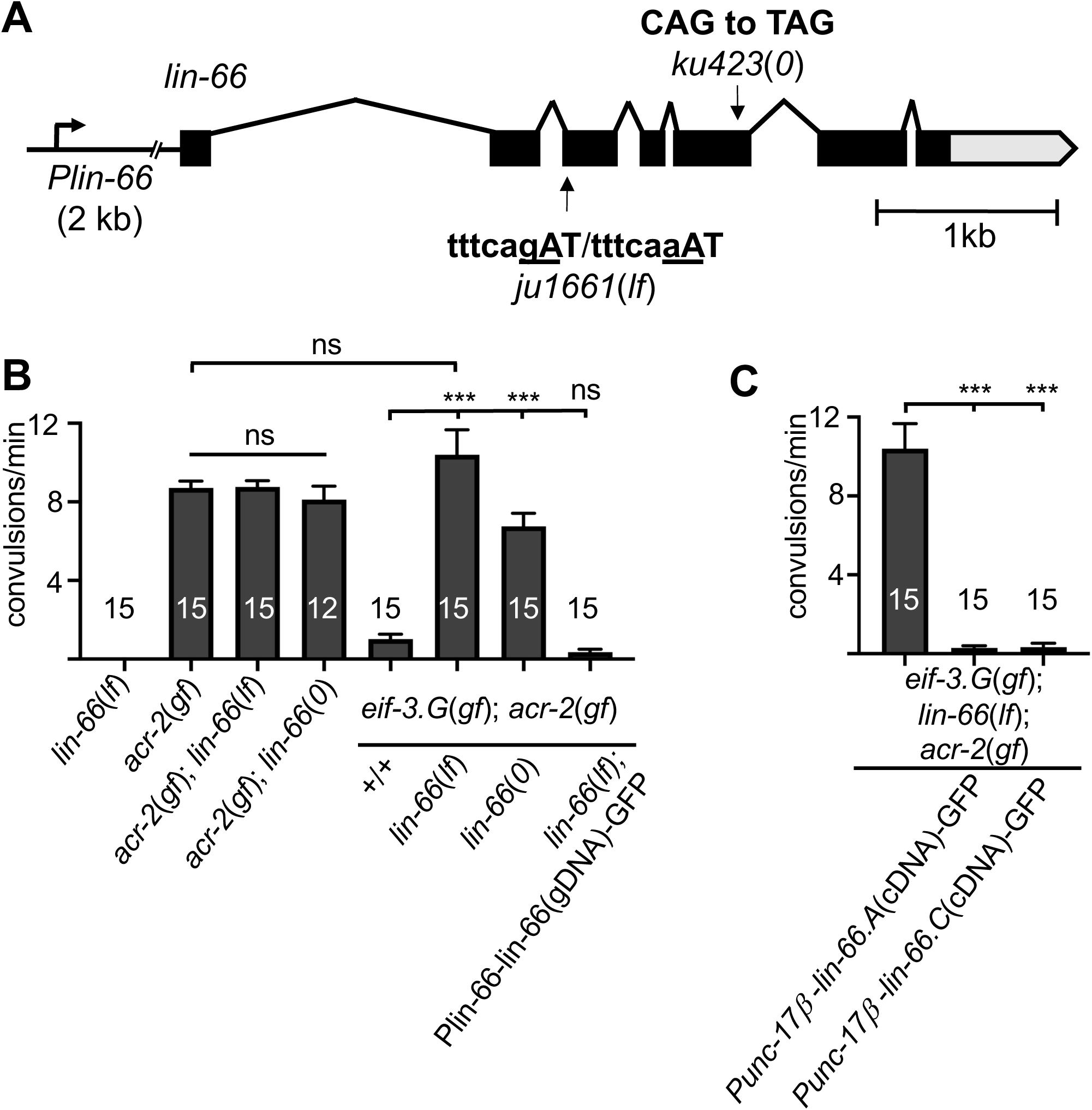
*lin-66* functions in cholinergic motor neurons to modulate *eif-3.g(gf)* activity. A) Illustration of the *lin-66* gene model on chromosome IV (modified from wormbase), along with the position and nucleotide change in *ju1661* and *ku423,* respectively. Black boxes represent exons, lines introns, gray box for 3’ UTR; and the promoter, *Plin-66*, includes 2 kb upstream sequences. B and C) Quantification of convulsion behavior in day one adult animals of the indicated genotypes. *lin-66*(*0*) animals were examined at L4 due to their larval arrest. Alleles used are: *eif-3.g(ju807gf), acr-2(n2420gf), ju1661* as *lin-66(lf), ku423* as *lin-66(0).* Convulsion quantification was from 2-3 independent observation, and at least two transgenic lines were scored. Statistics: one-way ANOVA with Bonferroni’s post-hoc test, (***) P<0.01, (ns) not significant.

The *lin-66* gene has seven exons and produces three mRNA isoforms (Figure 1A). The isoform a and c differ by 9 nucleotides as the result of alternative splice site usage at intron 3 and exon 4, and encode proteins of 627 aa and 624 aa, respectively. The isoform b has only exons 6 and 7, likely using alternative 5’ upstream sequences. To verify the effects of *ju1661* mutation, we isolated mRNAs from *ju1661* animals and obtained multiple *lin-66* cDNA fragments using primers annealing to exons 2 and 3. Sanger-sequencing analysis of these clones revealed both mis-spliced mRNA products using cryptic splice acceptor sites, which would result in out-of-frame beginning in the third exon, and also correctly spliced mRNA products as observed in mature mRNAs of wild type *lin-66*. Thus, *lin-66(ju1661)* causes partial defective pre-mRNA splicing and allows some production of wild-type proteins.

Null (0) mutations of *lin-66* were previously identified for their roles in temporal regulation of vulva cell fate (Morita and Han, 2006). *lin-66(ku423)* is a nonsense mutation changing amino acid Gln378, and causes developmental arrest at late larvae (L4 stage) (Figure 1A). By contrast, *lin-66(ju1661)* animals developed to fertile adults and displayed no discernable abnormalities in body shape, movement, and growth rate, consistent with the conclusion that *lin-66(ju1661)* retains partial function, designated as *lin-66(lf)*. *lin-66(lf); acr-2(gf)* double mutants showed convulsions indistinguishable from *acr-2(gf).* In *eif-3.G(gf); acr-2(gf)* background, *lin-66(lf)* restored convulsions to the level similar to *acr-2(gf)* single mutants (Figure 1B). We observed similar suppression effect in L4 animals of *lin-66(0); eif-3.G(gf); acr-2(gf)*. However, adult escapers of *lin-66(0); eif-3.G(gf); acr-2(gf)* showed less suppression effects, likely reflecting overall unhealthiness of *lin-66(0)*. We transgenically expressed wild type *lin-66* genomic DNA, containing 2 kb upstream sequences, the full coding region and 657 nt 3’ untranslated sequences, and observed complete reversal of the suppression effect associated with *lin-66(lf)* (Figure 1B). Together, these data indicate that loss of function in *lin-66* specifically reduces the gain-of-function effect of *eif-3.G(gf)* on *acr-2(gf)* convulsion behavior.

### LIN-66 acts in cholinergic motor neurons to mediate EIF-3.G(gf) activity on protein translation

*eif-3.G* and *lin-66* are broadly expressed in many types of cells (Blazie et al., 2021; Morita and Han, 2006). We previously showed that the suppression of *acr-2(gf)* convulsion by *eif-3.g(gf)* is due to a cell autonomous activity of *eif-3.g(gf)* in the cholinergic motor neurons. To test the cell type requirement of *lin-66,* we expressed full-length cDNAs of *lin-66.A* or *lin-66.C* specifically in cholinergic motor neurons using the *Punc-17b* promoter. We found that *lin-66(lf); eif-3.G(gf); acr-2(gf)* animals carrying such transgenes exhibited wild type locomotion resembling *eif-3.G(gf); acr-2(gf)* (Figure 1C). These data indicate that *lin-66* acts in the cholinergic motor neurons to interfere with *eif-3.G(gf)* function.

We previously reported that EIF-3.G(gf) selectively modulates protein translation efficiency on mRNAs with GC-rich 5’ UTRs to suppress convulsions of *acr-2(gf)* (Blazie et al., 2021). We next asked whether the genetic interaction between *lin-66* and *eif-3.G(gf)* involves regulation of protein translation by examining the expression of an EIF-3.G target, HLH-30. In *acr-2(gf)* single mutants the fluorescence intensity of HLH-30::GFP from a fosmid reporter (*wgIs433*) in the ventral cord cholinergic motor neurons was selectively and significantly elevated, and in *eif-3.G(gf); acr-2(gf)* double mutants, HLH-30::GFP expression in these neurons was reduced to levels in wild type animals (Figure 2) (Blazie et al., 2021). We found that in *lin-66(lf)* single mutants HLH-30::GFP intensity in the cholinergic motor neurons was similar to that in wild type animals. However, in *lin-66(lf); eif-3.G(gf); acr-2(gf)* animals, HLH-30::GFP expression was increased to the level comparable to that in *acr-2(gf)* single mutants (Figure 2). Additionally, *lin-66(lf)* did not further increase HLH-30::GFP expression in *acr-2(gf)* single mutants (Figure 2). Therefore, these data support a conclusion that *lin-66* mediates the activity of *eif-3.G(gf)* on protein translation.

**Figure 2.**
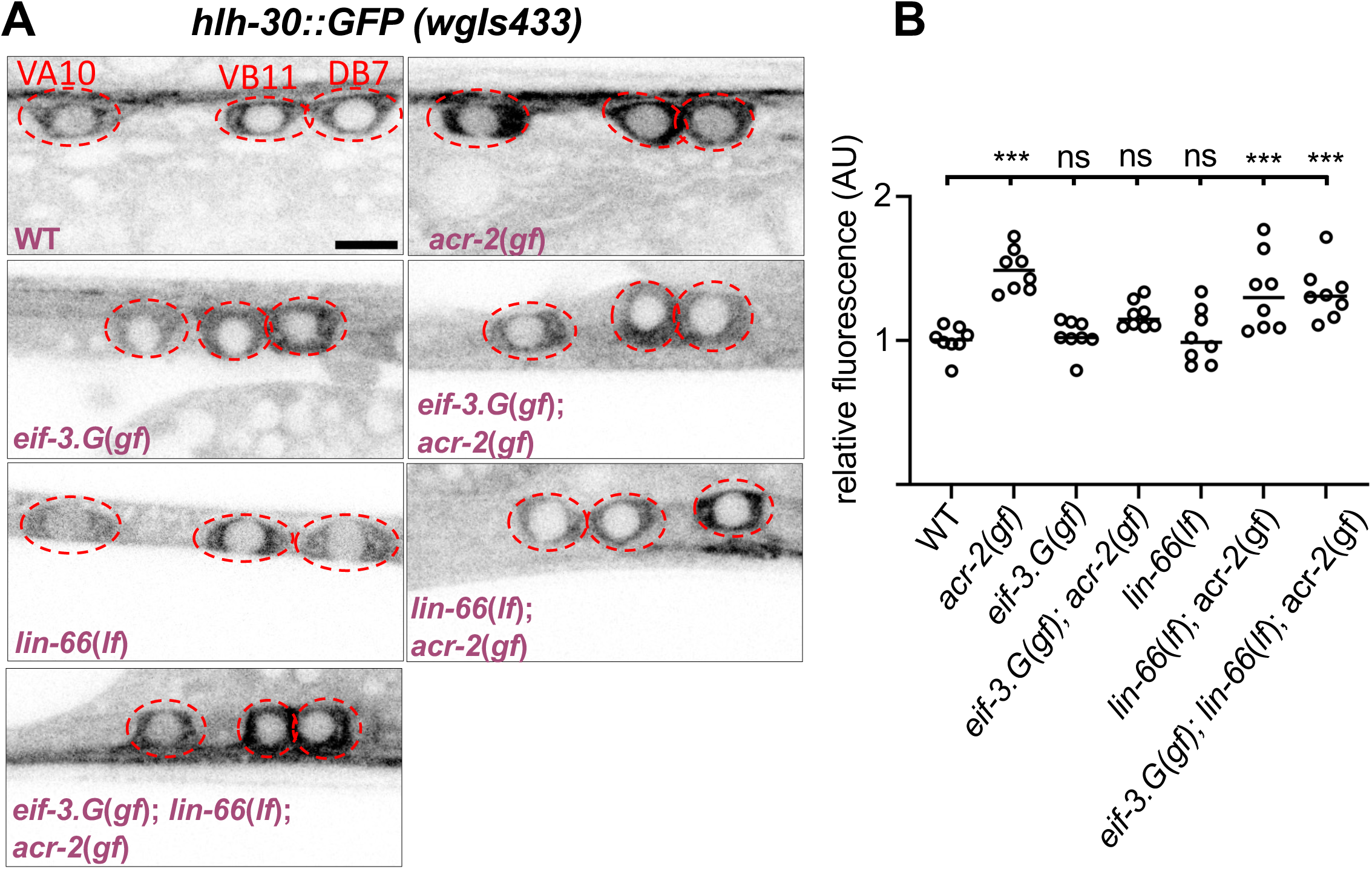
*lin-66* modulates protein translation in cholinergic motor neurons, dependent on EIF-3.G(gf) and ACR-2(gf). A) Representative confocal images of the cholinergic motor neuron soma from animals expressing *hlh-30::GFP* (a fosmid-based reporter, *wgIs432*). Cholinergic motor neurons were identified based on their stereotypic position, aided by a *Pacr-2-mcherry* reporter; the soma were enclosed by dotted red line; scale 4 μm. Alleles used are the same as in Figure 1A. B) Quantification of HLH-30::GFP intensity from each genotype (n= 8). Each dot represents the average fluorescence intensity from the VA10, VB11, and DB7 neuronal soma normalized to the wild type (WT) control. Statistics: (***) P< 0.001, (ns) not significant, one-way ANOVA with Bonferroni’s post hoc test.

### Expression of LIN-66 and EIF-3.G is largely independent of each other

We considered two possible explanations for how loss of *lin-66* function may reduce the activity of *eif-3.G(gf).* One is that *eif-3.G(gf)* might cause increased expression of LIN-66; alternatively, *eif-3.G(gf)* expression is dependent on *lin-66*. We first examined a previously published transgenic overexpression reporter line of *lin-66,* and observed comparable expression pattern in wild type and *eif-3.G(gf).* To more precisely determine LIN-66 expression, we used CRISPR-editing technology to tag the endogenously expressed LIN-66 with GFP fused in-frame at the C-terminus, allele designated *ju2002* (see Methods). However, the fluorescence intensity of the knock-in (KI) LIN-66::GFP was nearly invisible, indicating that LIN-66 is not abundantly expressed. We noticed that an intermediate GFP knock-in product that retained the *let-858* 3’ UTR used in the knock-in vector (Figure 3A, allele designated *ju1934*) showed detectable fluorescence in muscle, epidermis, and neuronal tissues throughout all developmental stages. Despite the increased expression, *lin-66::GFP(ju1934)* behaved as wild type, as homozygous *lin-66::GFP(ju1934)* animals were healthy and did not show any suppression effects on *eif-3.G(gf); acr-2(gf).* In the ventral cord motor neurons, LIN-66::GFP fluorescence was mostly concentrated within somatic cytoplasm. The overall intensity and subcellular pattern of *lin-66(ju1934)* was not altered in the presence of *eif-3.G(gf),* or in *eif-3.G(gf); acr-2(gf)* double mutants (Figure 3B). These observations argue against the idea that EIF-3.G(gf) regulates translation of LIN-66. Moreover, *lin-66(ju1934); acr-2(gf)* animals showed convulsion similar to *acr-2(gf),* suggesting that simply increasing LIN-66 expression is not sufficient to mimic the activity of EIF-3.G(gf). Conversely, we tested whether *lin-66* might regulate the expression of EIF-3.G. We found that functional GFP::EIF-3.G expressed from an integrated single-copy transgene showed similar intensity and localization pattern in wild type and *lin-66(ju1661).* To further address if the subcellular localization pattern of EIF-3.G and LIN-66 might depend on each other, we co-expressed an integrated single-copy transgene of *Peif-3.G-mKate2::eif-3.G(WT)* with *lin-66::GFP(ju1934)*. By confocal imaging analysis, we observed that the two proteins showed largely overlapping granular pattern and that the colocalization pattern of LIN-66::GFP and mKate2::EIF-3.G remained similar in both wild type and *acr-2(gf)* animals (Figure 3C). Based on these data, we conclude that LIN-66 and EIF-3.G do not regulate each other’s expression.

**Figure 3.**
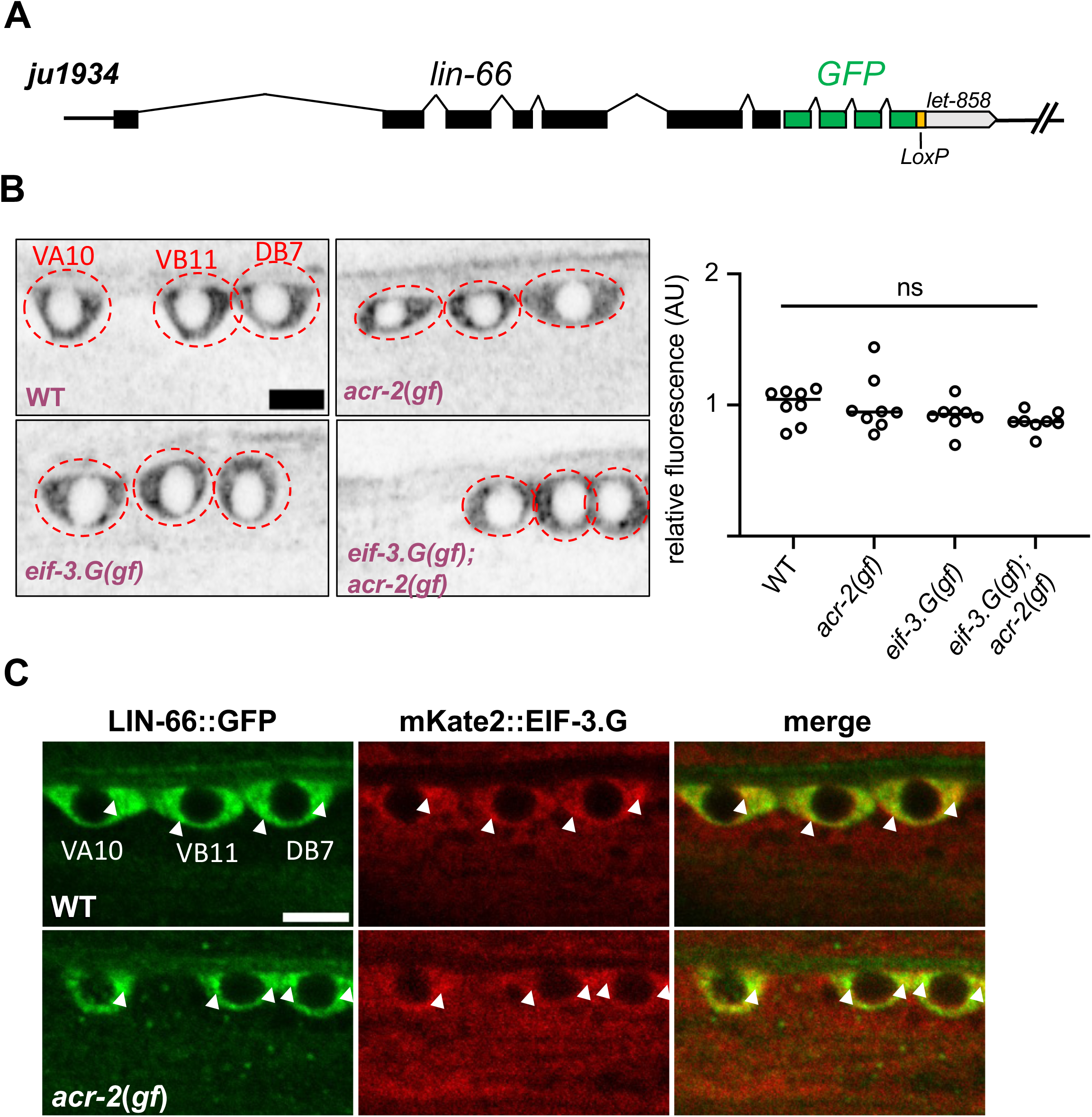
LIN-66 localizes predominantly to somatic cytoplasm and its expression pattern is not affected by *eif-3.g(gf)* or *acr-2(gf)*. A) *ju1934* has GFP knock-in to the C-terminus of *lin-66*, and retains the *let-858* 3’UTR followed by the self-excision cassette flanked by LoxP sites. B) Representative confocal images of LIN-66::GFP*(ju1934)* in the indicated cholinergic motor neuron soma (left) and quantification of GFP intensity in the three neuronal soma in each genotype (right, n=8 animals, scale 4 μm). Statistics, One-way ANOVA with Bonferroni’s post-hoc test; ns, not significant. Alleles used are the same as in Figure 1A. C) Representative confocal images of LIN-66::GFP and mKate2::*eif-3.G* expression in indicated cholinergic motor neuron soma; white arrows point to their colocalization in wild type and in *acr-2(gf);* scale 4 μm.

### LIN-66 localization and function depend on both a predicted structural region and low-complexity sequences

LIN-66 was previously reported to exhibit homology across nematode clades but with no apparent mammalian orthologs or identifiable domains. We therefore employed *in silico* tools to further analyze the LIN-66 protein sequence (see Methods). SMART protein domain analysis revealed several low complexity amino acid sequences at the N- and C-terminal region of LIN-66 (Figure S1), which was also predicted to be highly intrinsically disordered by PONDR. Additional prediction of folded protein structures using AlphaFold suggested amino acids 101-370 in the middle of LIN-66 to be a structured region (Figure S1, Figure 4A).

**Figure 4.**
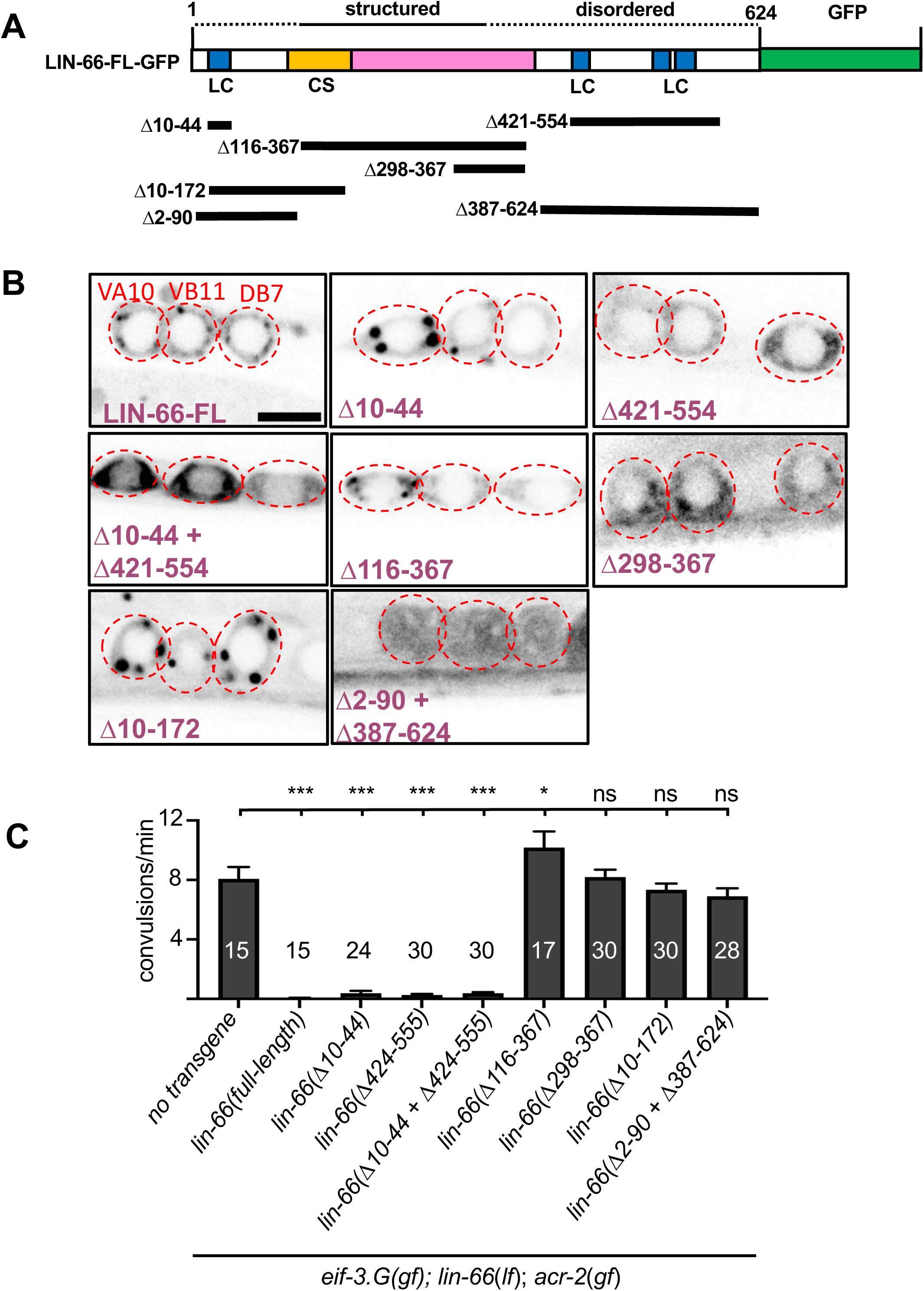
LIN-66 function depends on both the predicted structured region and flanking low complexity and intrinsic disordered sequences. A) Illustration of the LIN-66 protein (isoform C) tagged with GFP at the C-terminus. The colored domains include LC for low complexity sequence and CS for cold-shock domain. Dotted lines above represent predicted disordered regions and solid line for the structured region predicted by AlphaFold. Black bars below depict sequences deleted in *lin-66* truncation transgenes. B) Representative images of the indicated cholinergic motor neuron soma expressing each *lin-66*-GFP transgene illustrated in A: FL, full length; Δ indicates the amino acids deleted; scale 4 μm. C) Quantification of convulsion in animals expressing the *lin-66* transgenes shown in panel A-B. At least two transgenic lines for each expression construct were analyzed Statistics; One-way ANOVA with Bonferroni’s post-hoc test. ***, P<0.01; *, P<0.05; ns, not significant.

To test the functional relevance of the prediction, we generated a series of LIN-66 translation reporters using lin-66.C cDNA fused to GFP at the C-terminus and driven by the Punc-17β promoter, which is active only in the cholinergic motor neurons (Figure 4A). Overexpressed full length LIN-66::GFP showed somatic cytoplasmic localization with punctate pattern (Figure 4B). Removing the N-terminal disordered sequences, LIN-66(Δ10-44), did not change the overall localization pattern. Removing low complexity sequences at the C-terminus, LIN-66(Δ421-555), caused the protein to appear more diffuse (Figure 4B). Removing both N- and C-terminal low complexity sequences, LIN-66(Δ10-44 + Δ421-555), showed slightly increased expression but remained diffuse like LIN-66(Δ421-555) (Figure 4B). In *lin-66(lf); eif-3.g*(*gf*); *acr-2*(*gf*) triple mutant animals, all three LIN-66 mutant constructs showed rescuing activity, similar to the full length LIN-66 transgene (Figure 4C). In contrast, a transgene containing only the predicted structured region, LIN-66(Δ2-90, Δ387-624), showed a diffuse localization pattern and did not exhibit rescuing activity (Figure 4B-C). Other LIN-66::GFP transgenes that omitted some of the predicted structural region, LIN-66(Δ116-367), LIN-66(Δ298-367), and LIN-66(Δ10-172), did not affect the overall localization pattern, and also had no rescue activity (Figure 4B-C). These data indicate that the regions outside of the predicted structured region can influence LIN-66 punctate pattern, and that the predicted structured region is necessary, but insufficient, for LIN-66 function in *eif-3.g*(*gf*) mediated behavioral suppression of *acr-2(gf)*.

### LIN-66 function depends on a cold-shock domain

To further identify functional homologous domains of the structured region in LIN-66, we inputted the AlphaFold LIN-66 structure to the ProFunc server, which predicts protein function from structure (Laskowski et al., 2005). This analysis revealed that amino acids 101-170 are predicted to fold into a cold shock domain (Figure 5A), most similar to those present in *C. elegans* LIN-28 and human LIN-28 and CSDE1 (Figure 5B and Figure S2) (Jacquemin-Sablon et al., 1994; Moss et al., 1997). The observation that LIN-66(Δ10-172)::GFP did not show rescuing activity in *lin-66(lf); eif-3.g(gf), acr-2(gf)* (Figure 4B-C) supported the functional importance of the predicted cold shock domain.

**Figure 5.**
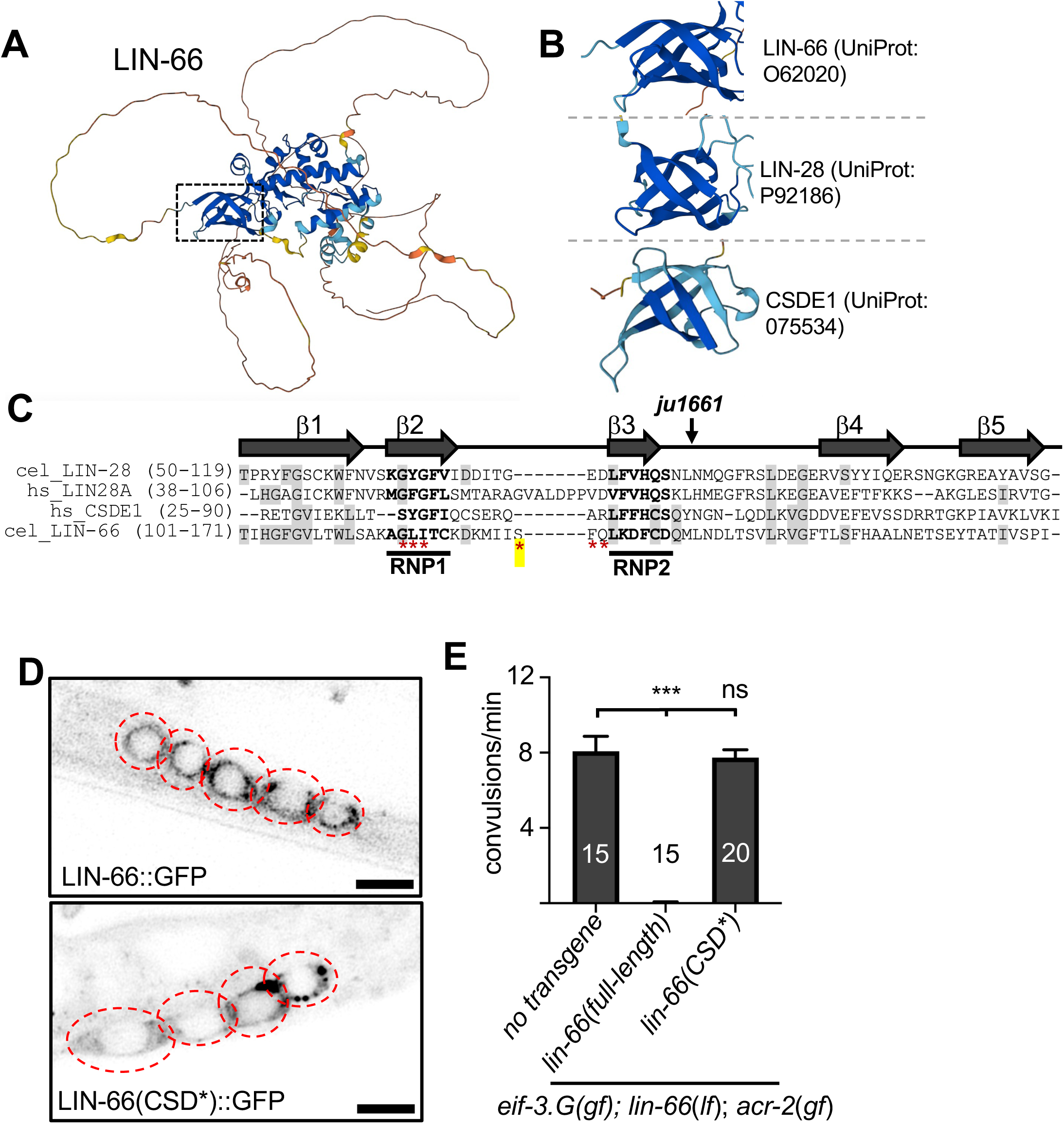
The cold-shock domain in LIN-66 is important for its function. A) The predicted structure of LIN-66 protein (UniProt accession: O62020) generated by the AlphaFold algorithm (https://alphafold.ebi.ac.uk/), with the box pointing to the cold-shock domain predicted by ProFunc analysis (Laskowski et al., 2005). B) The predicted LIN-66 cold shock domain resembles that in *C. elegans* LIN-28 and the first cold-shock domain of human CSDE1. C) Amino acid sequence alignment of cold-shock domains of *C. elegans* LIN-66 and LIN-28, and human LIN-28 and CSDE1. Bolded residues in β strands 2 and 3 indicate the RNP1 (R/K)-G-(**F**/**Y**)-(G/A)-(**F/Y**)-V-X-(F/Y) and RNP2 (L/I)-(**F**/**Y**)-(V/I)-X-(N/G)-L consensus sequences. Red asterisks represent the amino acids mutated to alanine or serine in the *lin-66(CSD*)* transgene. *ju1661* generates transcripts that would lead to LIN-66 out-of-frame beginning at the indicated amino acid within the CSD. D) Representative confocal image of the indicated cholinergic motor neuron soma (red dotted outline) expressing wild type *lin-66 (full-length)::GFP* or mutant *lin-66(CSD*)::GFP* transgene; scale 4 μm. E) Quantification of convulsion from animals with indicated genotype and transgenes. At least two transgenic lines were scored. Statistics, One-Way ANOVA with Bonferroni’s post-hoc test; ***, p<0.01; ns, not significant.

Cold shock domains generally consist of five β-strands, with the second and third β- strands containing highly conserved amino acids necessary for RNA binding (Heinemann and Roske, 2021). By amino acid sequence alignment, the predicted cold shock domain of LIN-66 showed significant homology in the conserved regions, notably the conserved glycine in the β1 strand and the conserved amino acid residues within the RNP1 and RNP2 motifs (Figure 5C). To test whether the residues in RNP motifs are important for LIN-66 function, we mutated them in the full-length LIN-66::GFP translation reporter, designated *lin-66(CSD*).* When expressed in the cholinergic motor neurons, we observed that LIN-66(CSD*)::GFP exhibited a diffuse pattern with prominent puncta in neuronal soma (Figure 5D), similar to that observed with full-length LIN-66. However, the *lin-66(CSD*)* transgene did not show any rescuing activity in *lin-66(lf); eif-3.G(gf); acr-2(gf)* triple mutants (Figure 5E). Taken together, these data show that the predicted cold-shock like domain in LIN-66 is required for its function in EIF-3.g(gf)-dependent regulation of neuronal hyperexcitation, likely involving its RNA-binding activity.

## Discussion

The eIF3 complex possesses a variety of functions to modulate protein synthesis by engaging the unique activities of its thirteen subunits (Cate, 2017). We previously showed that *C. elegans* EIF-3.G conveys a specific regulation of protein translation via the conserved zinc finger to modulate neuronal activity of the cholinergic motor neurons (Blazie et al., 2021). We now provide evidence that this specialized activity of EIF-3.G requires LIN-66, a protein previously described to regulate protein translation through an unknown mechanism during animal development (Morita and Han, 2006). Furthermore, by combining structural informatic analysis with *in vivo* functional dissection, we have uncovered a cold-shock domain in LIN-66 that is critical for its function. We propose that LIN-66 acts as a cell-type and context-dependent facilitator to mediate the activity of EIF-3.G in protein translation.

*C. elegans lin-66* was discovered in a genetic screen for genes regulating vulva cell fate. Null mutants of *lin-66* arrest in late-stage larvae, indicating that it has essential roles in animal development (Morita and Han, 2006). *lin-66* exhibits complex genetic interactions with multiple genes known to regulate temporal cell fate specification. Moreover, in *lin-66* null mutants the expression of the cold-shock domain protein LIN-28 was increased, which may partly involve the 3’ UTR of *lin-28* mRNA. The authors proposed that *lin-66* may act in a post-transcriptional regulatory network to modulate protein translation, although they speculated that *lin-66* might not directly act on *lin-28* mRNA. In our *acr-2(gf)* induced cholinergic overexcitation paradigm, through unbiased genetic screening, we were able to isolate a partial loss of function mutation of *lin-66* that displays no discernable developmental defects, but specifically ameliorates the gain-of-function effect of EIF-3.G on locomotion behavior and on protein translation in cholinergic motor neurons. Thus far, we have not yet found any evidence supporting a role of *lin-28* in *acr-2(gf)* induced neuronal overexcitation. Our transgenic expression studies support a conclusion that the genetic interaction between *lin-66* and *eif-3.g* is specific to the cholinergic neuron hyperexcitation. We also show that LIN-66 mediates protein translation modulation through EIF-3.G targets that has GC-rich 5’ UTR. LIN-66 and EIF-3.G co-localize to the same subcellular domains in motor neuron soma, and do not appear to regulate each other’s expression. We thus hypothesize that the two proteins provide functional co-dependence for regulation of protein translation.

Little is known about the function of LIN-66, which is likely due to both the lack of identifiable protein domains and the low abundance of LIN-66 protein ((Morita and Han, 2006), and this study). Through *in silico* analyses of available structural and informatic database, we found that LIN-66 is predicted to have a structured region embedded within low-complexity sequences. Within the predicted structured region, we further identified a cold-shock domain. Cold-shock domains were originally named because of their presence in multiple bacterial proteins that were induced by cold shock (Heinemann and Roske, 2021; Lindquist and Mertens, 2018; Mihailovich et al., 2010; Sommerville, 1999). It is now known that cold shock domains are present in many proteins with diverse functions, independent of any response to cold shock. For example, human Y-box binding protein 1 participates in transcription, splicing and translation (Kleene, 2018). The LIN-28 proteins use a combination of their cold-shock domain with CCHC zinc fingers to regulate microRNA metabolism or processing (Moss and Tang, 2003). Other cold-shock domain-containing proteins can act as RNA chaperones or facilitate RNA processing or degradation (Lindquist and Mertens, 2018). Interestingly, several protein translation factors, such as eIF1A, eIF2α or eIF5A, all contain a cold-shock domain (Amir et al., 2018), although the function of these cold-shock domain remains undefined.

Our *in vivo* structure-function dissection studies support the predicted LIN-66 protein structure. We find that mutating the conserved residues in the RNP motifs abrogates LIN-66 function, without altering the overall protein expression, suggesting that LIN-66 has the potential to directly bind nucleic acids. Full-length LIN-66 localizes to somatic cytoplasm with visibly detectable puncta or granules. We find that low-complexity sequences of LIN-66 contribute to the punctate pattern of LIN-66. LIN-66 function depends on both the structured region and some low-complexity sequences. Low complexity sequences are generally intrinsically disordered, and many studies have shown that proteins with intrinsically disordered regions, most notably those involved in forming RNA granules, have the propensity to undergo protein phase separation (Lee et al., 2022). Conceivably, LIN-66 function may engage multi-valent types of interactions to modulate protein synthesis by co-opting translation machinery in specialized cellular contexts.

Dysregulation of protein translation is associated with many human neurological disorders (Kapur et al., 2017). Altered expression of eIF3g is linked to narcolepsy and autism (Choi and An, 2021; Holm et al., 2015). Cold-shock domain-containing proteins are also broadly associated with RNA metabolism and protein translation (Lindquist and Mertens, 2018). Through unbiased forward genetic screening for locomotion behaviors associated with altered neuronal circuit activity, we have uncovered a highly selective protein translation regulatory network involving the interaction between the conserved EIF-3.G subunit and a cold-shock domain-containing protein LIN-66. Our findings underscore the power of forward genetic analysis in deciphering functional specificity underlying how an essential component of protein translation complex can tune the neuronal proteome to balance neural circuit outputs.

## Acknowledgements

We thank our lab members for valuable discussions throughout the work, M. Han for providing some of the *lin-66* reagents. Some strains were provided by the CGC, which is funded by NIH Office of Research Infrastructure Programs (P40 OD010440). SMB and DF were trainees on a T32 training grant (NS007220). This work is supported by NS R37 035546 and NS R35 127314 (to YJ).

## Author contribution

S. M. B. and Y. J. conceived the project, S. M. B. performed the genetic screen and carried out and supervised experiments; D. F. contributed to imaging analysis; Y. Z. assisted in genetic mapping of *lin-66(ju1661)* and RT-PCR analysis. S. M. B. and Y. J. wrote the manuscript, with inputs from D.F. and Y. Z.

## Materials and methods

### C. elegans genetics

All *C. elegans* strains were maintained at 20°C on Nematode Growth Media (NGM) seeded with *E. coil* OP50 as described (Brenner, 1974). Behavior and imaging analysis was performed in young adult or L4 hermaphrodites; males were used for crosses. Compound mutants were generated by genetic crossing using standard procedures. Mutant alleles were detected using a combination of visual observations of locomotion and/or fluorescence reporters, and further verified by identification of allele-specific DNA changes using PCR and sequencing. All strains used in this study are listed in Table S1. Information of mutant alleles and the primers for PCR amplification is listed in Table S2.

### Genetic screening for suppressors of *eif-3.G(gf)*

We conducted a clonal genetic suppressor screen using CZ21759 *eif-3.G(ju807gf); acr-2(n2420gf)* double mutants, following the standard procedure (Brenner, 1974). Briefly, mix-staged *eif-3.G(gf); acr-2(gf)* double mutants were washed into a 2 mL solution of M9 media with 47 mM ethyl methanesulfonate (Sigma) and incubated at 22°C on a spinning wheel for 4 hours. Worms were pelleted in a Beckman Allegra X-14R centrifuge at 1,500 rpm for 2 minutes and washed twice with M9 media, with centrifugations in between washes. Worms were transferred to a fresh seeded NGM plate and allowed to recover for 20 minutes at 22°C. Two mutagenized L4 (P0) animals were picked to 17 plates and cultured at 20°C. After three days, eighteen F1 animals from each P0 (total 306 F1s) were singly transferred to freshly seeded NGM plates, and cultured at 20°C for 2 more days. Young adult F2 animals on each F1 plate were screened for any restored convulsion behavior by visual inspection under a dissection microscope. Many initial isolates did not produce viable progeny, and seven F2 animals produced true-breeding lines, but with variable convulsions resembling *acr-2*(*gf*) single mutants. We outcrossed the original isolates using N2 males and were able to re-isolate one suppressor, *ju1661,* that showed consistent suppression and that was dependent on both *eif-3.g(gf)* and *acr-2(gf)*.

### Mapping of *lin-66*(*ju1661*)

Genomic DNAs were purified from strains CZ26710 and CZ26711 using Gentra Puregene Tissue Kit (Qiagen). Whole genome DNA library preparation and 90 bp paired-end sequencing was performed by Beijing Genomics Institute (BGI, USA). Sequencing reads of 20X coverage were mapped to the *C. elegans* reference genome (ce10) using Burrows-Wheeler Aligner (BWA) (Li and Durbin, 2009). Genome Analysis Tookid (GATK) (McKenna et al., 2010) was used to locate single-nucleotide polymorphisms in the mapped reads comparing to the reference genome. The mapped SNPs were annotated by their relative proximity to the nearest gene model (WS220) using SnpEff version 2.1a (Cingolani et al., 2012). We used custom SQL scripts to identify SNPs that are detected in both the original isolate (CZ26710) and the 1x outcross (CZ26711) and excluded from CZ21759 (sequences reported in (Blazie et al., 2021)). We performed further outcrossing to N2 and re-isolated multiple *ju1661* recombinants. Using the mutagenesis-induced SNPs to perform fine mapping of these recombinants, we located *ju1661* to a small interval of chromosome IV containing *lin-66*.

### CRISPR-editing to knock in GFP to *lin-66*

We used the CRISPR-Cas9 SEC (self-excision-cassette) method (Dickinson et al., 2013) to insert GFP in-frame immediately preceding the stop codon in the endogenous *lin-66* locus. Homology arm sequences that included 500 bp upstream (primers YJ12796 and YJ12797) and 500 bp downstream (primers YJ12794 and YJ12795) of the *lin-66* stop codon were PCR amplified from N2 genomic DNA. Silent mutations were introduced in the portion of the homology arm containing the sgRNA sequence to prevent Cas9 cleavage of the repair template. Gibson Assembly (NEB) was performed to ligate the resulting PCR amplicons with the pDD268 vector (Addegene #132523) digested with AvrII and SpeI, producing the clone pCZ1053. The sgRNA sequence 5’-CGTTTGAACTCACTCCGTAT -3’ was incorporated into pDD162 vector to produce pCZ1054, following the protocol as described. A DNA mix containing 10 ng/μL pCZ1053, 50 ng/μL pCZ1054, 10 ng/μL pGH8, 5 ng/μL pCFJ104, and 2.5 ng/μL pCFJ90 was injected into N2 young adult hermaphrodites. The P0 animals were cultured at 25°C for 3 days, and their progeny were treated with 250 ug/μL hygromycin. After another three days, surviving progeny that showed the roller phenotype but did not express co-injection mCherry markers were selected. A single line, designated *ju1934,* was confirmed by PCR genotyping to contain GFP in-frame to the last amino acid of LIN-66. To remove SEC, ∼16 CZ29347 *lin-66*(*ju1934*) L1 animals were transferred to a freshly seeded NGM plate and heat shocked at 34°C for 6 hours. After culture at 22°C for 3 days, animals without the rolling phenotype were selected and verified for the presence of GFP and absence of the SEC, resulting in an allele designated *ju2002*.

### Molecular cloning and transgenic expression of *lin-66* wild type and variants

Full-length *lin-66* genomic DNA including 2 kb upstream sequences and 650 nt downstream of the stop codon was obtained by PCR amplification from wild type (N2) genomic DNA and cloned into Gateway entry vector to generate pCZGY3547. cDNA for *lin-66A* and *lin-66C* were obtained from wild type cDNA library and cloned into Gateway entry vector to generate pCZGY3561 and pCZGY3560, respectively. GFP fusion expression constructs were made by Gibson Assembly using primers listed in Table 2. All truncated (Δ) *Plin-66*::*lin-66C*::GFP expression constructs were generated using the phosphorylated primers listed in Table 2. Briefly, primer pairs for each Δ DNA fragment were used to PCR amplify a portion of pCZGY3555 (Plin-66-lin-66C::GFP) and the product was treated with DpnI to remove the template and DpnI deactivated at 80°C for 25 minutes. The resulting product was then subjected to intramolecular ligation using T4 DNA ligase and transformed into DH5a. All final clones were verified by Sanger sequencing and listed in Table 2. A mixture of 20 ng/μL transgene and 70 ng/μL 1 kb ladder (NEB) filler DNA was microinjected into CZ26711 young adult hermaphrodites, following standard procedure (Mello et al., 1991). More than two transgenic lines were selected by screening F1 progeny of microinjected animals for GFP expression and transmission efficiency.

### *in silico* annotation of LIN-66 protein structure

We used the simple modular architecture tool (http://smart.embl-heidelberg.de/; (Letunic et al., 2015)) and SEG (https://mendel.imp.ac.at/METHODS/seg.server.html;(Wootton, 1994)) to locate low complexity (LC) amino acid sequences in lin-66 isoform C. Predictors of Natural Disordered Regions (PONDR; http://www.pondr.com/; (Xue et al., 2010)) was used with default settings to assess disordered sequence regions of LIN-66. The structured region between amino acids 101-376 and cold-shock domain (amino acids 101-171) was identified from the output of AlphaFold software (https://alphafold.ebi.ac.uk/) using LIN-66 (UniProt: 062020) with default settings (Jumper et al., 2021). Structural homology between the cold-shock domain of LIN-66 and human CSDE1 was identified by inputting the entire LIN-66 isoform C amino acid sequence into the ProFunc database (http://www.ebi.ac.uk/thornton-srv/databases/profunc;(Laskowski et al., 2005)) using default settings.

### Quantification of convulsion behavior

Convulsion behavior was assessed by visual inspection of day one young adult hermaphrodites under a dissection microscope according to the following protocol. L4 animals were transferred to a fresh seeded NGM plate the day prior to behavioral assay. The following day, each single young adult animal was placed onto the bacterial lawn of an NGM plate seeded with OP50 and allowed to acclimate for 90 seconds. Convulsions, defined as a brief shortening of the animal body length accentuated by muscle contraction, were quantified over a 90 second period of continuous monitoring. Observers were blinded to genotype. Convulsions in *lin-66*(*0*) animals were quantified at L4 stage because their development arrests before reaching adult. Our data reports the average convulsion frequency over a 60 second interval.

### Confocal imaging and analysis

L4 or young adult animals were immobilized in a small droplet of 1mM levamisole in M9 liquid media on a microscope slide containing a 2% agarose pad. Imaging was performed with a Zeiss LSM800 confocal microscope using 63X lens and identical image settings (1.25 mm pixel size with 0.76 ms pixel time, 50 mm pin-hole), unless otherwise indicated. We used *juEx2045(Pacr-2-mCherry)* to identify the VA10, VB11, and DB7 cholinergic motor neurons as previously described (Blazie et al., 2021). All other imaged cell types were identified based on their stereotyped position and anatomical features. Observers were blinded to genotype when possible.

### Quantifying fluorescence in motor neuron soma

All quantification of fluorescence intensity was performed using the integrated density function in ImageJ (Schindelin et al., 2012). We acquired the mean integrated density from a selected region around each VA10, VB11, and DB7 soma and subtracted background from an equivalent area of each image. The resulting values were normalized to the mean fluorescence intensities obtained from the same GFP reporter in same neuronal soma in a wild type background.

### Statistical analysis

GraphPad Prism9 software was used for all statistical analysis and p-values below 0.05 were considered significant. Sample sizes were determined via power analysis as in our previous studies (Blazie et al., 2021; McCulloch et al., 2017; Takayanagi-Kiya et al., 2016).

---

**Figure S1:**
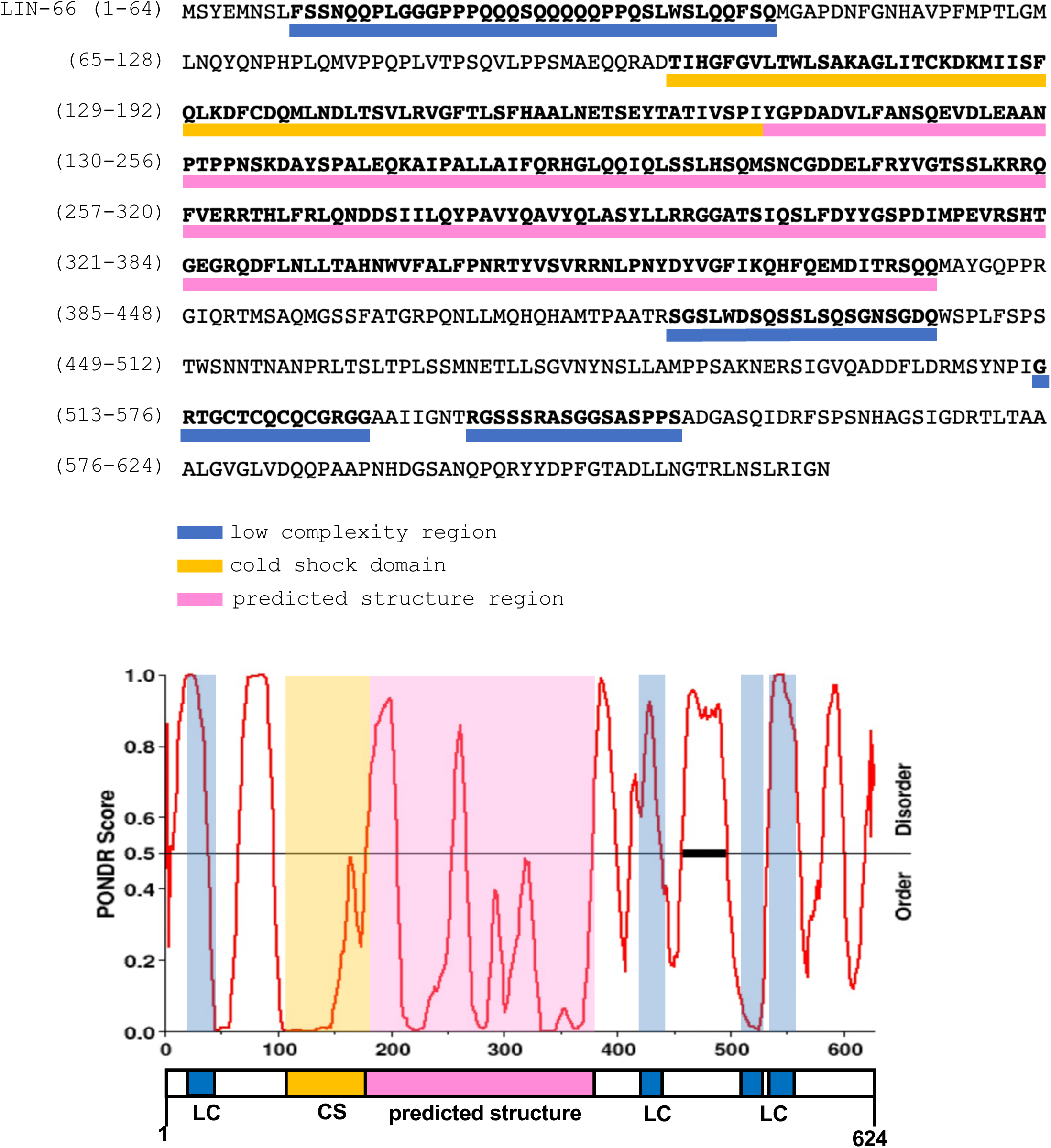
*in silico* sequence analysis of LIN-66. Top shows *C. elegans* LIN-66 isoform C protein sequence with indicated domains highlighted by colored bars below the corresponding amino acid sequence. Lower graph is PONDR analysis of LIN-66 disordered sequences. PONDR scores > 0.5 indicate relative disorder. The LIN-66 protein structure is displayed below the graph.

**Figure S2:**
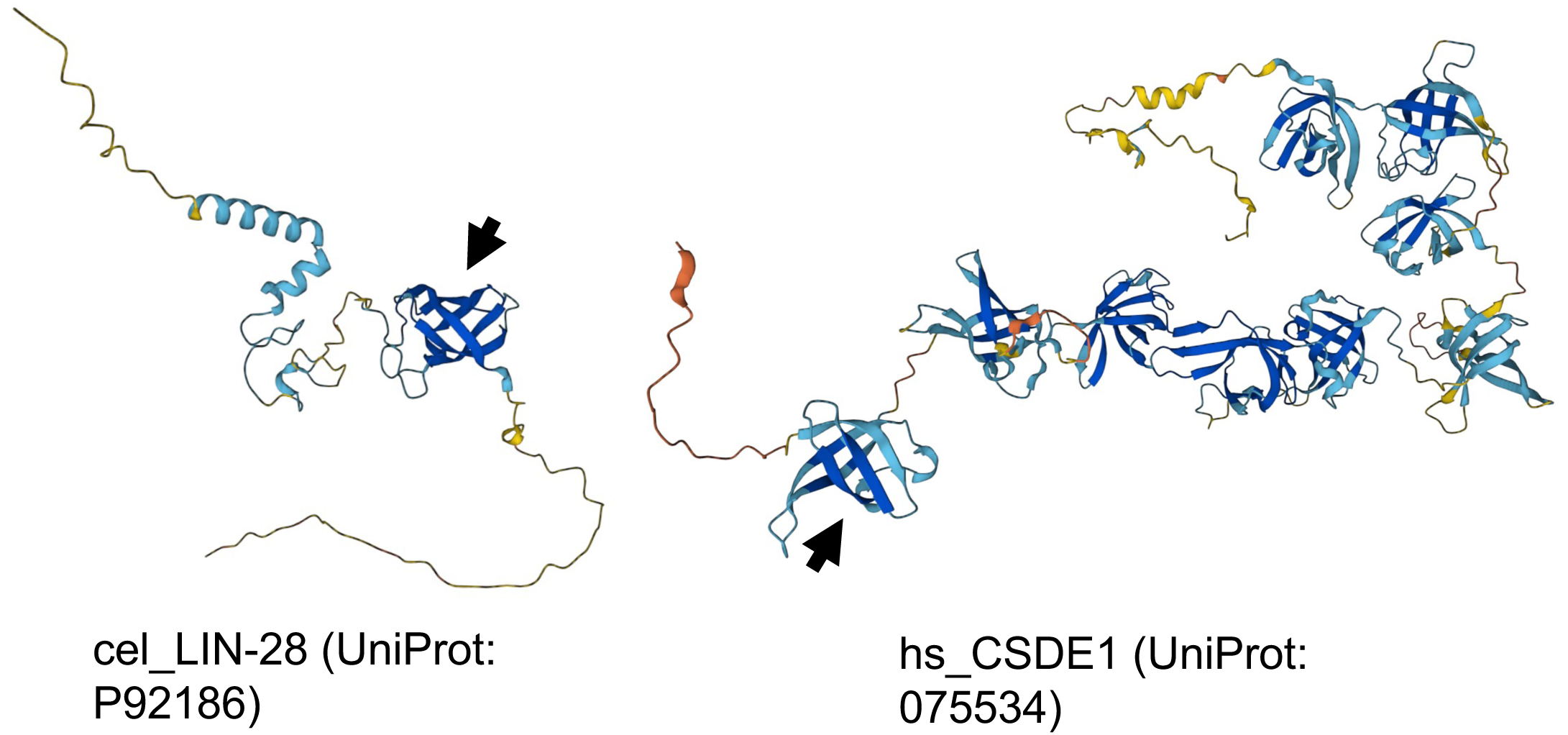
AlphaFold predicted structures of *C. elegans* LIN-28 and human CSDE1. Arrows indicate the cold shock domain with homology most similar to the cold shock domain revealed in *C. elegans* LIN-66.

**Table S1:**
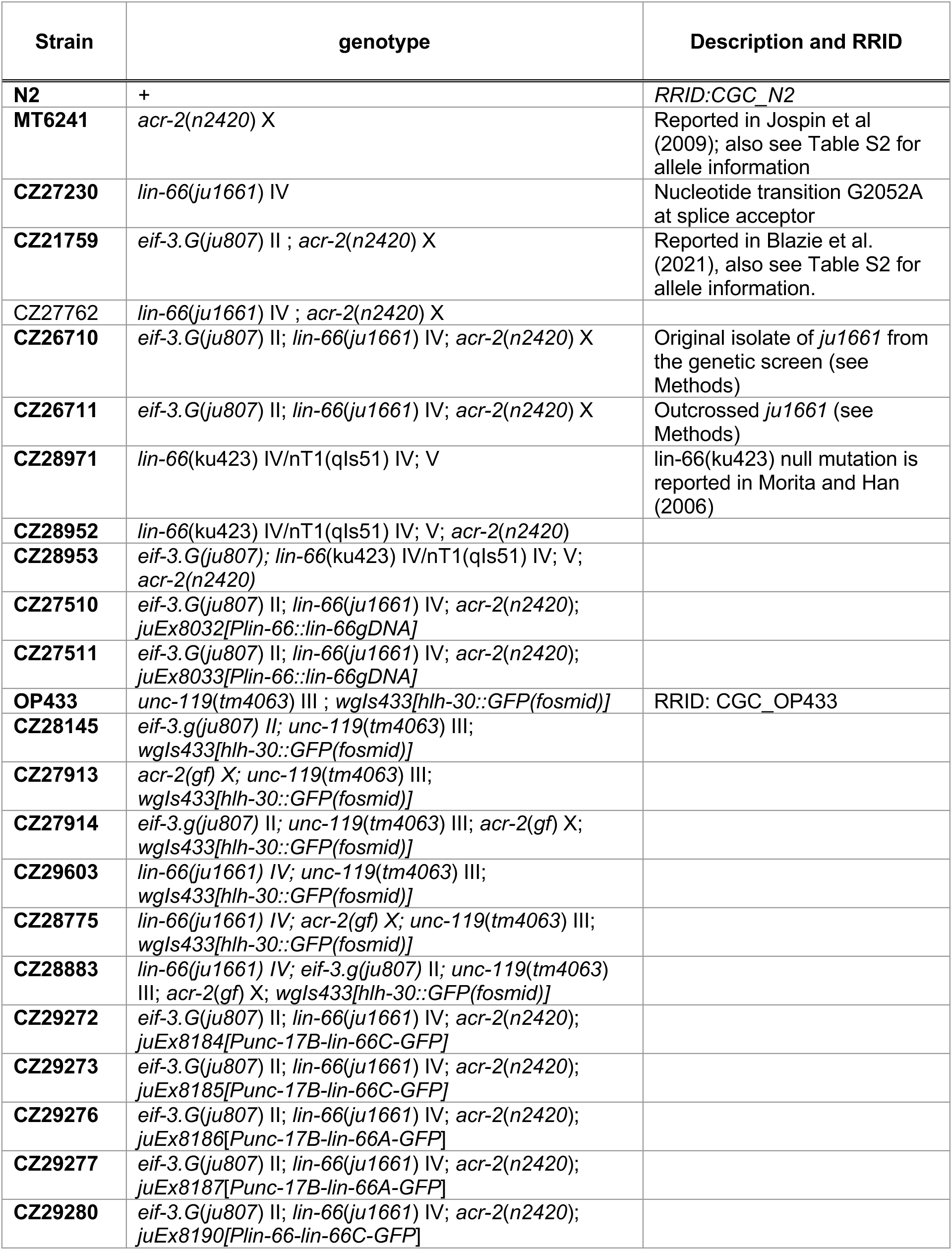
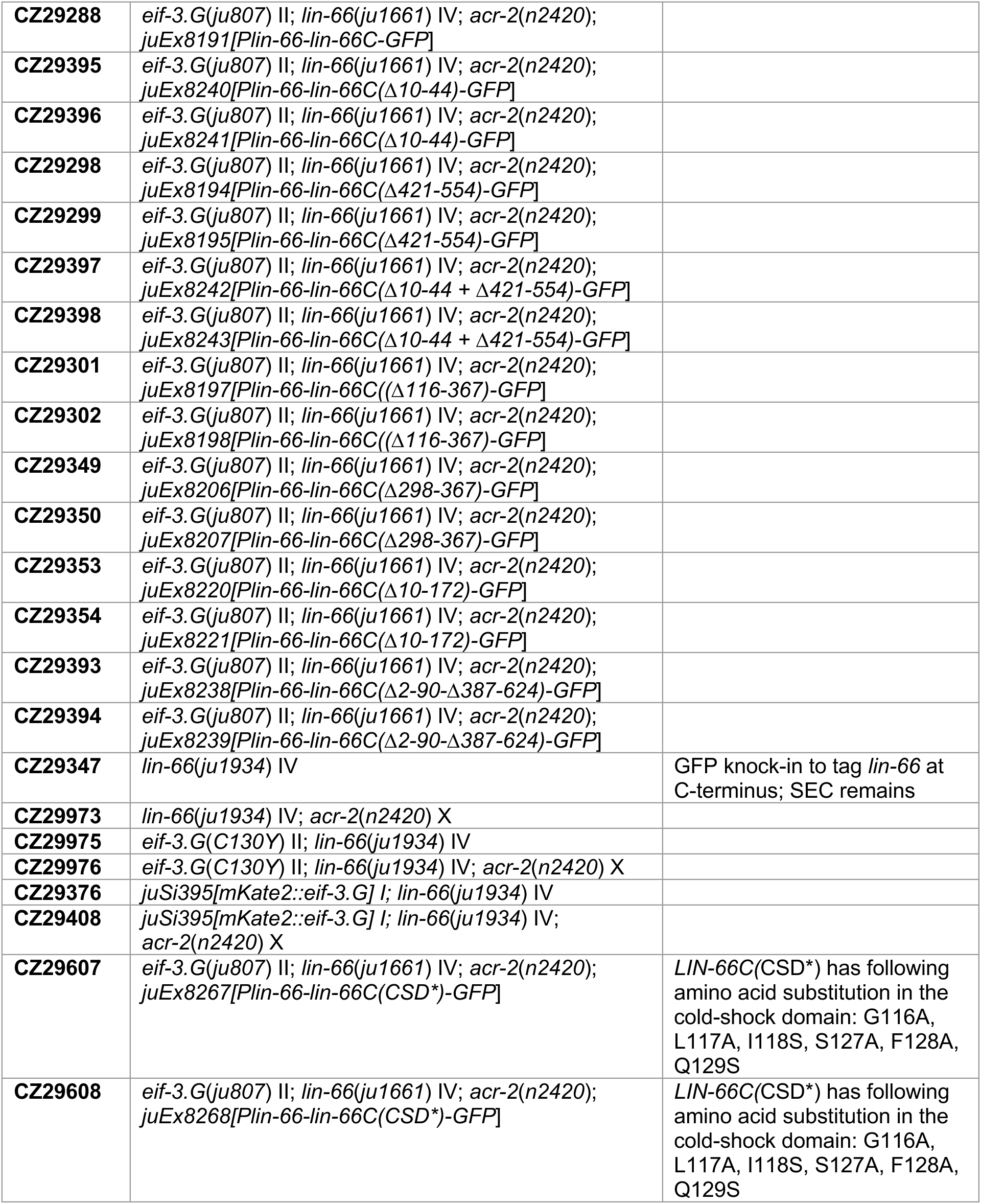
Strain and genotype information.

**Table S2:**
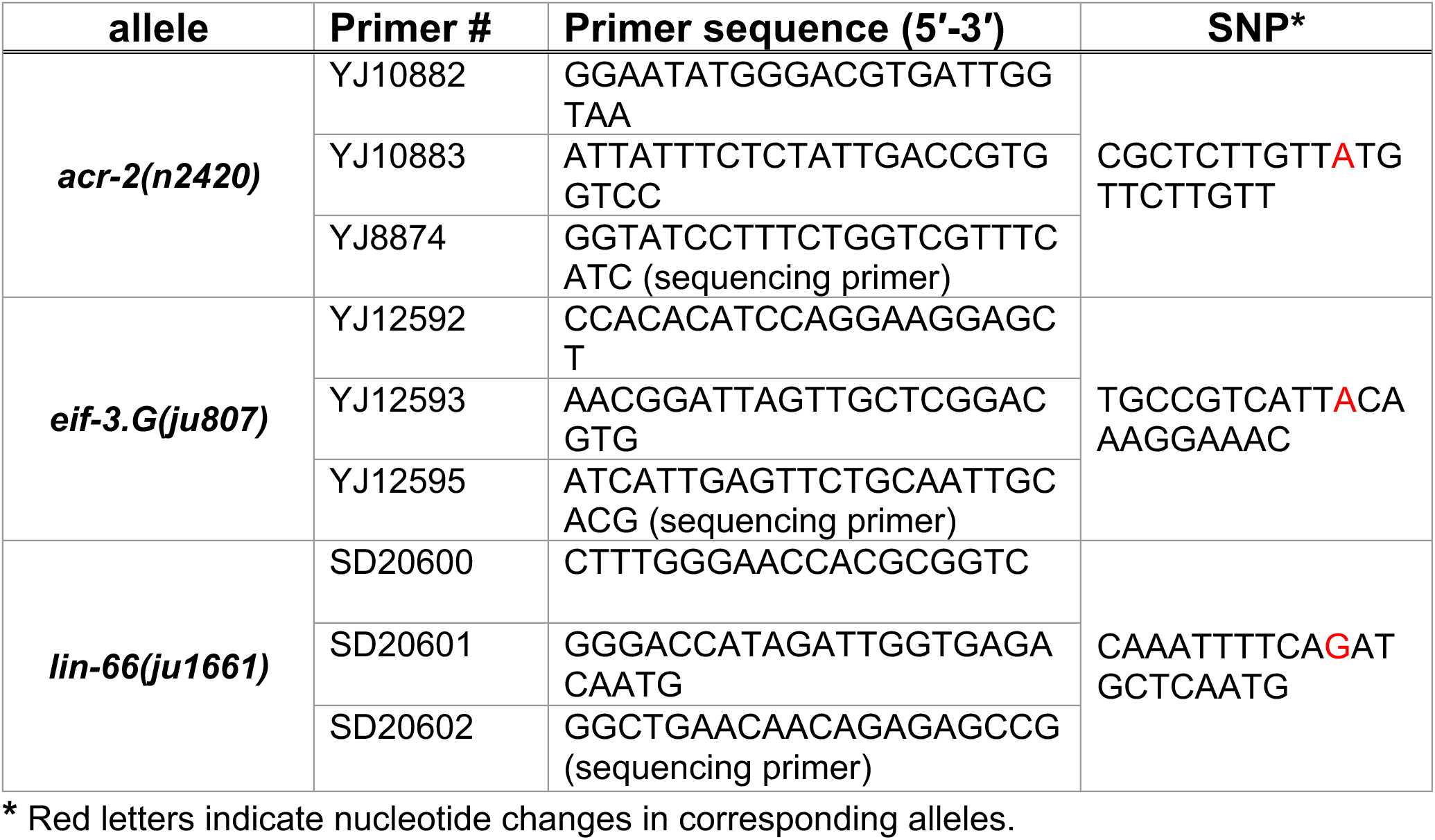
Allele Genotyping information.

**Table S3:**
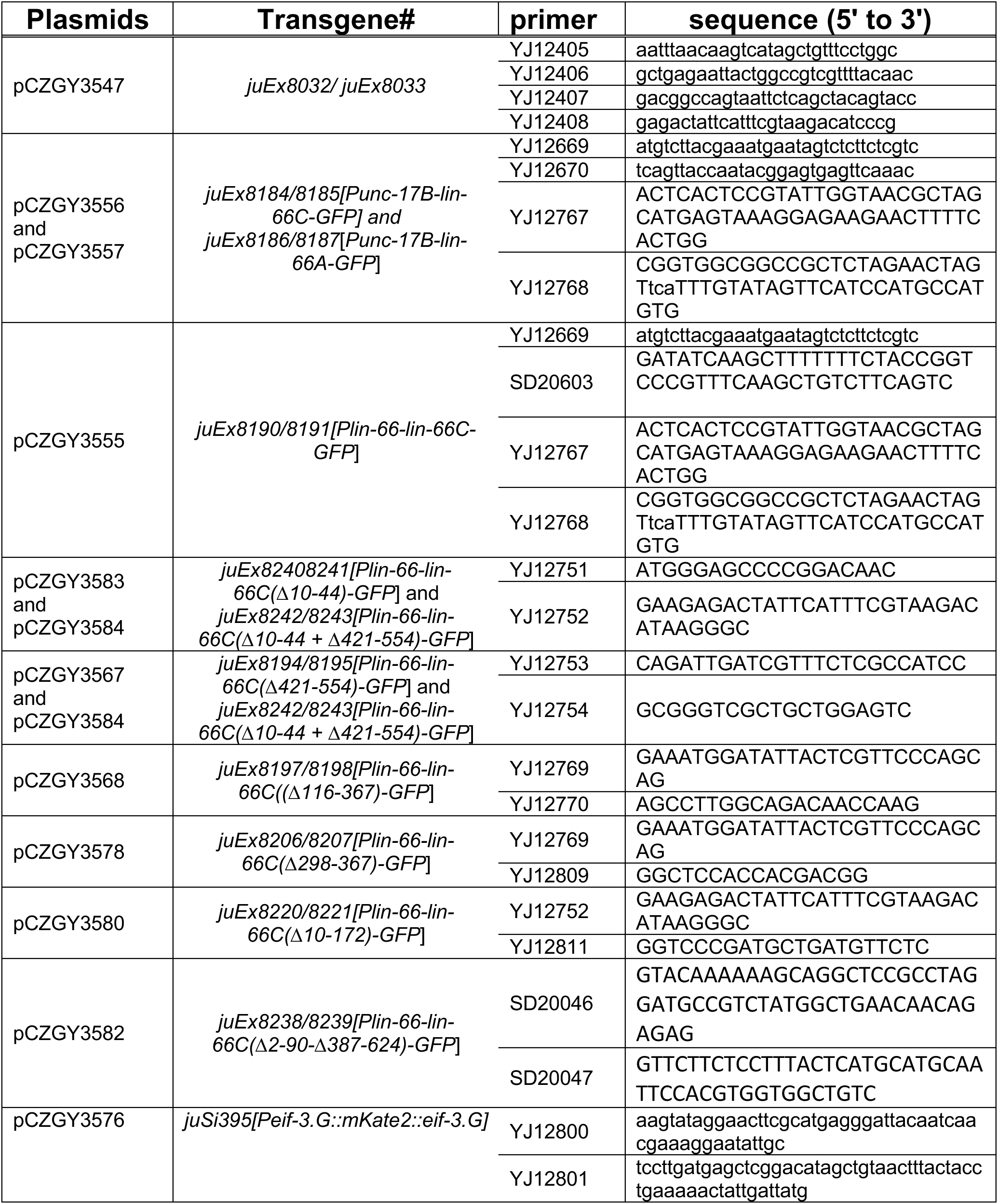
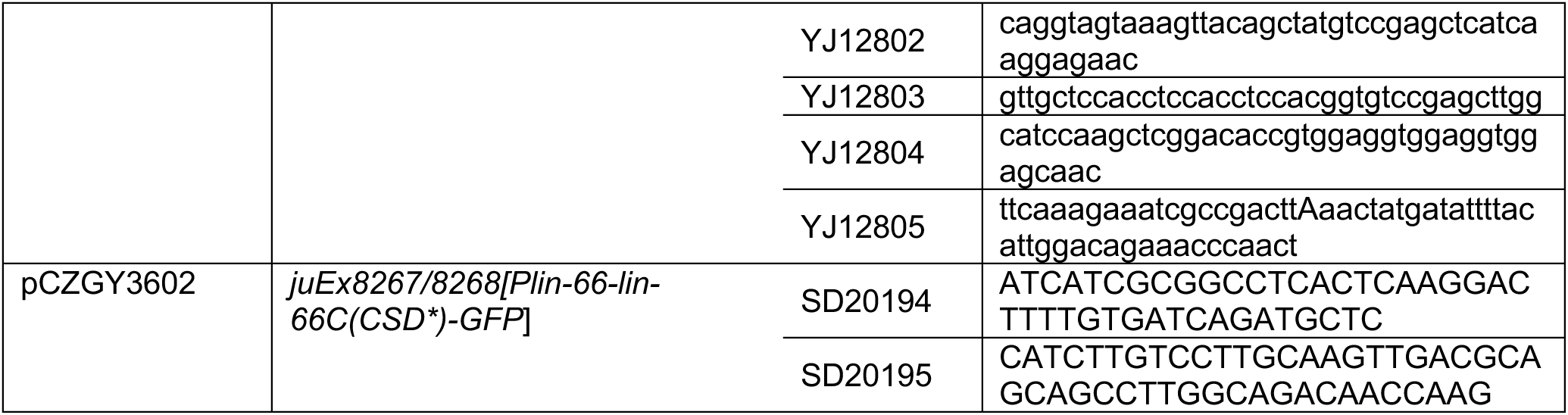
DNA plasmids and primers used for cloning.

